# Diet specialization in an insular population of coastal Peregrine Falcons

**DOI:** 10.1101/2021.09.07.459223

**Authors:** Luke J. Sutton, Sebastian W. Loram

## Abstract

Individual diet specialization is known to occur in populations of generalist predators, where specific individuals develop specialist feeding strategies. Diet specialization has been reported in many raptor species, and it may be an important driver of intraspecific population structure. Here, we quantify the diet of five breeding pairs of Peregrine Falcons *Falco peregrinus* from an offshore island determined from prey remains collected over four breeding seasons. Three prey species accounted for 69.8 % of total prey frequency, with Manx Shearwater *Puffinus puffinus* the primary prey accounting for 47.3 % by frequency and 40.8 % by biomass. Herring Gull *Larus argentatus* was the second most important prey species by frequency (13.8 %) and biomass (29.8 %) followed by Domestic Pigeon *Columba livia* (frequency = 8.7 %, biomass = 7.0 %). Predation frequency on specific prey groups varied substantially between breeding pairs and months. Two pairs specialized on Manx Shearwater, one pair specialized on Herring Gull and Manx Shearwater, with the remaining two pairs having a relatively generalist diet of Manx Shearwaters, Domestic Pigeon and small passerines. Predation on Manx Shearwaters increased throughout the breeding season with a peak in total diet frequency of 63.8 % in July, with a concurrent decrease in Herring Gull predation frequency. Higher percentage of Manx Shearwater in the diet was able to explain 87 % of the variation in a narrower dietary breadth for the Peregrine pairs. Our results suggest individual diet specialization may be important for understanding population density in insular raptor populations.

## Introduction

Ecological specialization is a key concept in ecology and evolutionary biology (Bolnick *et al*. 2003, Araujo *et al*. 2011), with individual diet specialization (IDS) a potential mechanism promoting coexistence between individuals through specialized use of food resources (Schoener 1974, Wiens 1984). Populations of diet generalist species are often comprised of a subset of diet specialized individuals (Pagani-Nunez *et al*. 2016). This may reduce resource-use overlap because specialized individuals utilize a narrower range of resources than conspecific competitors. IDS is expected to be common amongst species occupying higher trophic levels, where intraspecific competition for resources is expected to be high (Navarro-Lopez *et al*. 2014). Recently, studies of IDS within raptor populations have established this behaviour may be common in this bird group (e.g., Constantini *et al*. 2005, Whitfield *et al*. 2009, Navarro-Lopez *et al*. 2014, Terraube *et al*. 2014). Specialized individuals within raptor populations may have higher reproductive success than generalists (Dawson *et al*. 2011), though that assertion may not always apply (Whitfield *et al*. 2009). Therefore, understanding IDS is important because of its potential to regulate intraspecific population dynamics and breeding success (Bolnick *et al*. 2003, Ceia & Ramos 2015).

The Peregrine Falcon *Falco peregrinus* (hereafter ‘Peregrine’) is a widely distributed raptor with a global range, and considered a diet generalist, feeding on a broad range of other bird species (White *et al*. 2013). Though IDS has been described in several Peregrine populations from North and South America (Beebe 1960, Verlade 1993, Castellanos *et al*. 2006, Garcia *et al*. 2014), and anecdotally from the British Isles (Ratcliffe 1993), quantitative data is lacking from much of the Peregrine’s global range. In the British Isles, the Peregrine has a largely generalist diet (e.g., Dixon 2005, Sutton 2015a), with Domestic Pigeon *Columba livia* the primary prey (Ratcliffe 1993; Dixon *et al*. 2018). The British Isles has many small offshore islands off its western seaboard, some of which have dense sub-populations of breeding Peregrines relative to their area (Ratcliffe 1993; Sutton 2015b, Sutton 2016). Ratcliffe (1993) suggested that the population density of these island Peregrines may be driven by seabird abundance, which are likely to comprise the main food source for this insular Peregrine sub-populations.

Recent dietary assessments from Lundy, an island in the Bristol Channel between England and Wales, indicate that seabirds, including Manx Shearwater *Puffinus puffinus* and Herring Gull *Larus argentatus*, are the primary food source for the resident Peregrines (Sutton 2016; Sutton *et al*. 2017). Lundy was historically one of many important seabird colonies lying off the south west coast of the British Isles (Brown *et al*. 2011). Since the 1900’s seabird populations on the island have declined due to predation from invasive predators and human over-harvesting. After the eradication of Black *Rattus rattus* and Brown Rats *R. norvegicus* on the island in 2006 and formal legal protection, there has been an increase in breeding seabirds, in particular Manx Shearwater (Booker & Price 2014). Lundy also has a high breeding density of Peregrines, with five territorial pairs resident from 2002 to the present (Sutton 2016). The factors influencing this high breeding density are complex and not fully understood. However, one contributory factor may be diet specialization, allowing reduced competition for food resources, possibly enabling more pairs to breed in close proximity. Here, we assess foraging behaviour and diet specialization in this insular Peregrine population during the breeding season between the years 2014 and 2017. We aim to: (**1**) quantify diet composition and spatio-temporal patterns of predation, and (**2**) determine the level of individual diet specialization amongst the five resident breeding pairs.

## Methods

### Study area

Lundy (51°11’N, 04°40’W) is a small offshore island located in the Bristol Channel, England, 18 km from the nearest mainland point in the administrative county of Devon. The island is approximately 5-km long by 1-km wide, with 13-km of mainly granite sea cliffs. Elevation on the island plateau ranges from 80–143 m/asl, covering an area of approximately 445 ha. Two-thirds of the island plateau is comprised of unimproved grassland, moorland and maritime heath, dominated by Common Heather *Calluna vulgaris*. The southern third of the island supports a small tourist and farming community where the dominant land use is semi-improved and improved grassland.

### Data collection

We collected prey samples on a monthly basis between April and August 2014–2017 from five territorial breeding pairs. We followed a pre-determined circuit of the island recording food items from prey remains (including feathers, bones, and carcasses) left after feeding (Marti *et al*. 2007). We conducted systematic transect searches within territories, checking regular plucking areas along cliffs and slopes adjacent to nest cliffs. Prey remains were assigned to a territory if located within a 400-m radius of the nest cliff. This distance was determined as the minimum cross-over radius between territories to prevent recording outside of a known territory. We defined territories on cardinal points of the compass as: L1/NE (North-East); L2/E (East); L3/S (South); L4/SW (South-West); L5/W (West). Any remains found outside the 400-m radius from nest cliffs were assigned as random events and not used in the spatial analyses.

Prey remains were removed for identification and the location recorded with a handheld Global Positioning System (GPS) device to prevent double recording on subsequent visits. Remains observed on inaccessible feeding areas were recorded using a Digital Single Lens Reflex (DSLR) camera for later analysis and the location recorded indirectly using Google Earth (https://earth.google.com/web/) software. We identified prey species from Brown *et al*. (2003) and previous experience (Sutton 2015a, 2016; Sutton *et al*. 2017). Avian prey remains from Peregrines are readily identified. On carcasses the breast is ruptured with distinct notches in the breastbone from the Peregrines’ beak when feeding (Bang & Dahlstrom 2011). Plucked feathers will often be found concentrated in a pile surrounding the carcass indicating a kill, distinct from gull feeding remains (Corkhill 1973). Any raptor kills recorded during the breeding period were assumed to be from Peregrines as they are the only resident breeding raptor on Lundy (Davis & Jones 2007).

### Statistical analysis

We divided prey items into nine prey groups (Auks, Corvids, Ducks/Waders, Gulls, Lagomorphs, Pigeons/Doves, Shearwaters, Small Passerines, Other), and five weight classes (<50 g, 51–250 g, 251–500 g, 501–1000 g, >1000 g). We compared spatial and temporal variation in total prey frequency between territories and months using Fisher’s Exact test. We calculated dietary niche breadth (*B*) on the sum total of all prey items using the formula of Levins (Krebs 1999),

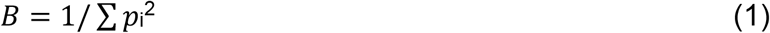

where *p*_i,_represents the proportion of the diet contributed by the ith taxon. Niche breadth was then standardized (*B*_*A*_) for easier interpretation following Hurlbert (1978),

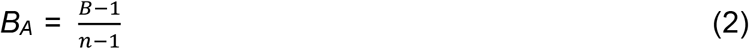

Values for this index range on a scale of 0.0 to 1.0 with lower values towards zero indicating narrower dietary breadth and higher values towards one indicating broader diet breadth. We used linear regression to model the relationship between prey weight and log-transformed frequency of predation, and to model the relationship between percentages of Manx Shearwater with total dietary breadth from each territorial pair. All statistical analyses were performed in R (v3.5.1, R Core Team 2018).

## Results

In total, we recorded 520 prey samples to a specific species or genus, with 36 prey species identified (Table S1, Appendix 1), comprising 35 birds and one mammal, European Rabbit *Oryctolagus cuniculus*. Avian prey species ranged in size from Common Chiffchaff *Phylloscopus collybita*, eight grams), up to Mallard *Anas platyrhynchos*, 1210 g). Three prey species comprised 69.8 % of total diet by frequency. Manx Shearwaters were the most common prey species by frequency (47.3 %) and biomass (40.8 %), followed by Herring Gull (frequency = 13.8 %, biomass = 29.8 %) and Domestic Pigeon (frequency = 8.7 %, biomass = 7.0 %). Prey items in the 251-500 g weight class were taken most frequently (62.1 %, Table 1), followed by the 500-1000 g weight class (18.9 %). The most common prey group were Shearwaters (comprised wholly of Manx Shearwater) by both frequency and biomass, followed by Gulls (Fig. 1). There was a positive relationship between prey mass and frequency of predation, with increased predation on prey species with greater mass (Fig. 2), explaining 16 % of the variation in prey frequency (*F*_*1,34*_ = 6.69, *R*^2^ = 0.16, *p =* 0.014).

**Table 1.** Percent frequency of predation by prey weight groups

| Prey weight group (g) | Frequency (%) |
| --- | --- |
| <50 | 11.5 |
| 51-250 | 3.7 |
| 251-500 | 62.1 |
| 501-1000 | 18.9 |
| >1000 | 3.9 |
| Total | 100.0 |

**Figure 1.**
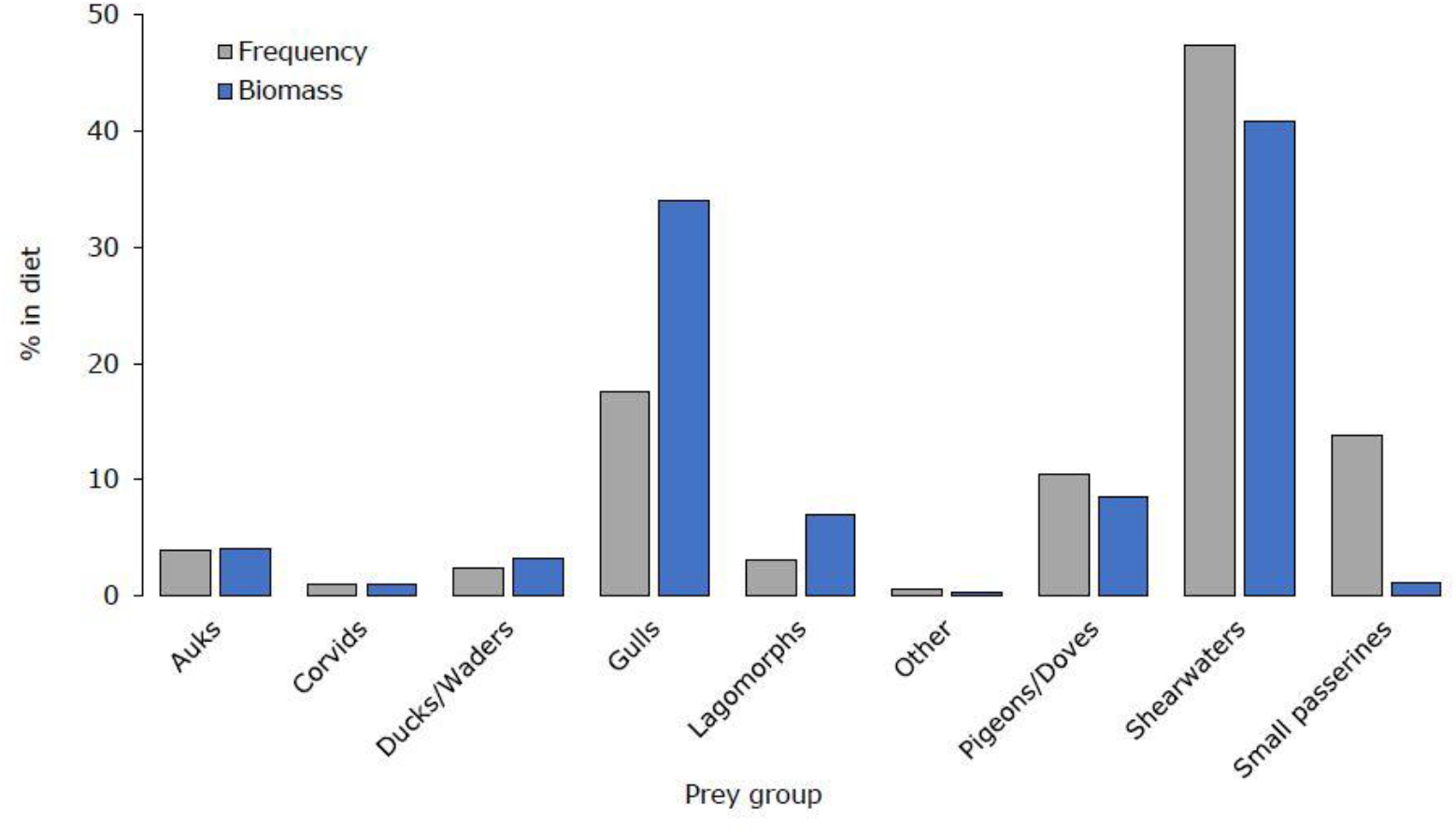
Prey groups by total frequency and biomass (%) predated by Peregrine Falcons breeding on Lundy from April-August 2014-2017.

**Figure 2.**
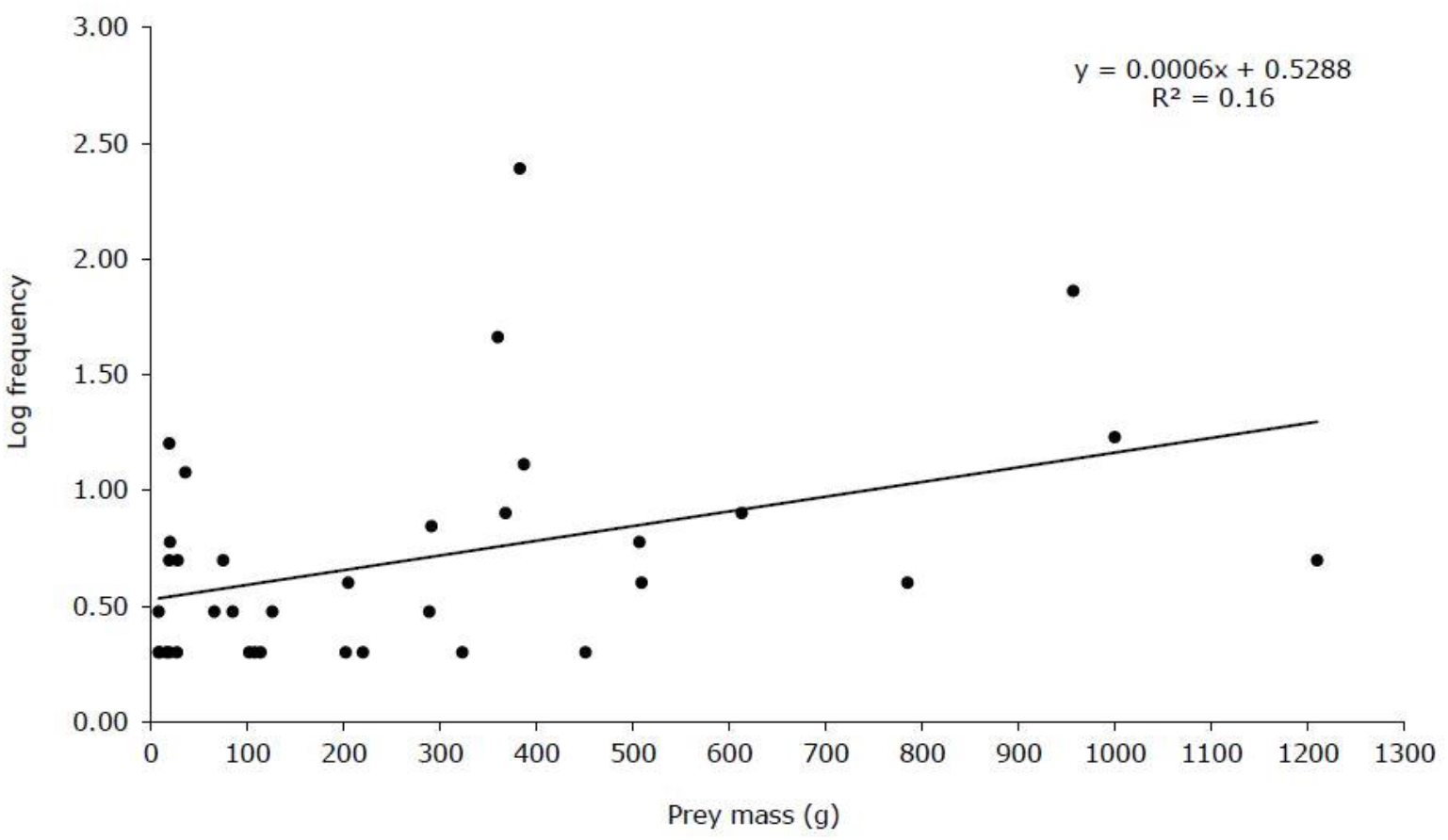
Predation frequency by mass of individual prey species taken by Peregrine Falcons breeding on Lundy from April-August 2014-2017. Unit mass (g) taken from BTO (2016).

Prey group diet composition varied substantially between nest territories (Fisher’s Exact = *p* <0.001, Fig. 3). Peregrine pairs in territories L1/NE and L4/SW had a high proportion of Shearwaters indicating a specialized diet (L1/NE = 53.7 %, L4/SW = 62.6 %). Pairs in territories L2/E and L3/S showed a generalist diet of Pigeons/Doves, Shearwaters and Small Passerines (Fig. 3). The pair in territory L5/W were diet specialists of both Shearwaters (47.1 %) and Gulls (35.0 %), with the pair in territory L4/SW also regularly taking Gulls, albeit less frequently (20.9 %). Frequency of prey group predation also varied substantially over the five-month (April-August) sampling period (Fisher’s Exact = *p* <0.001, Fig. 4). Relatively equal numbers of Gulls, Shearwaters and Small Passerines were predated in April, with a steady increase in total frequency of Shearwater predation from May onwards, peaking in July at 63.8 %. Conversely, as frequency of Shearwater prey group predation increased, Gull prey group predation frequency dropped to a low of 6.3 % by July, before a marked increase in August (26.3 %).

**Figure 3.**
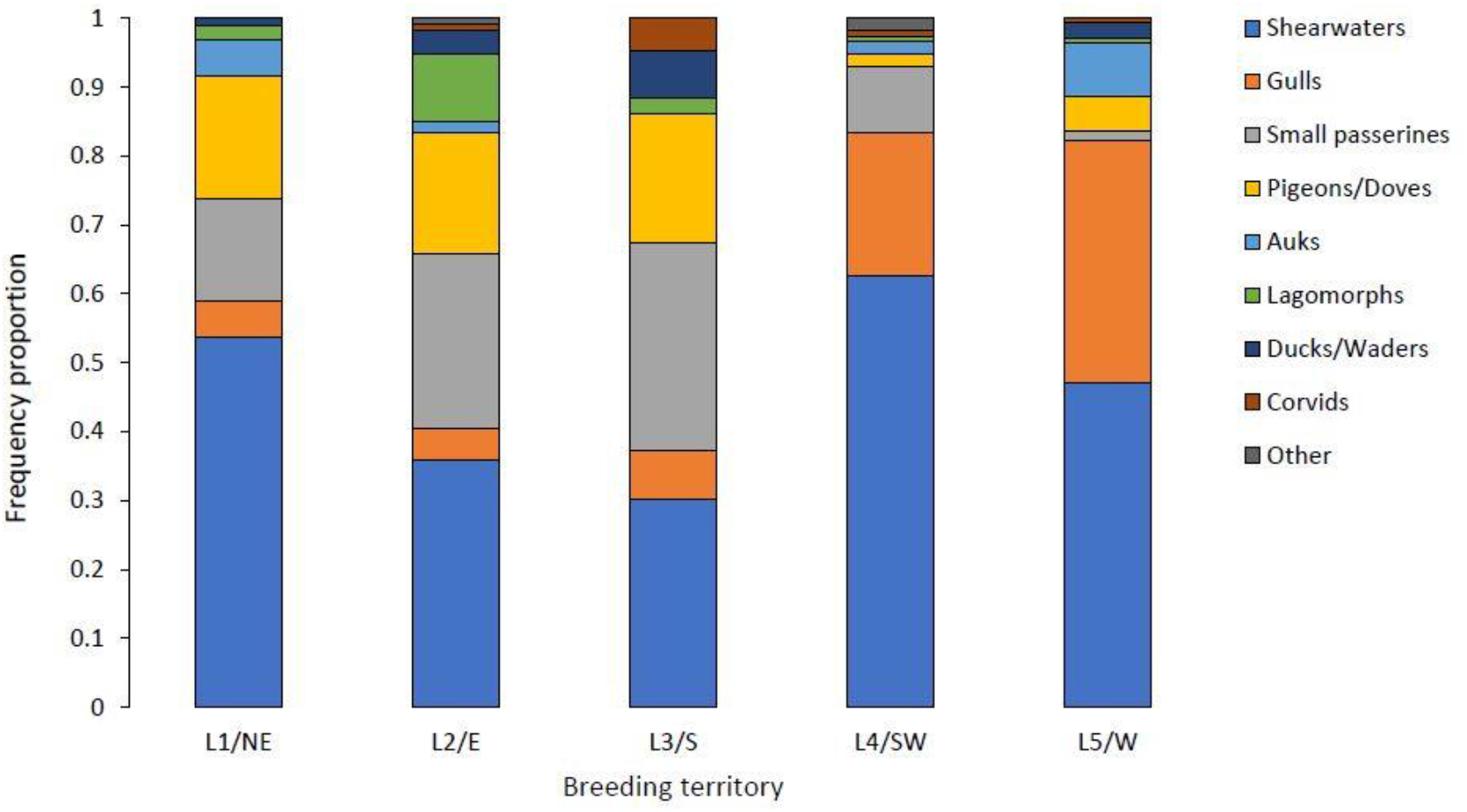
Spatial variation between nest territories in frequency of prey groups predated by Peregrine Falcons breeding on Lundy from April-August 2014-2017.

**Figure 4.**
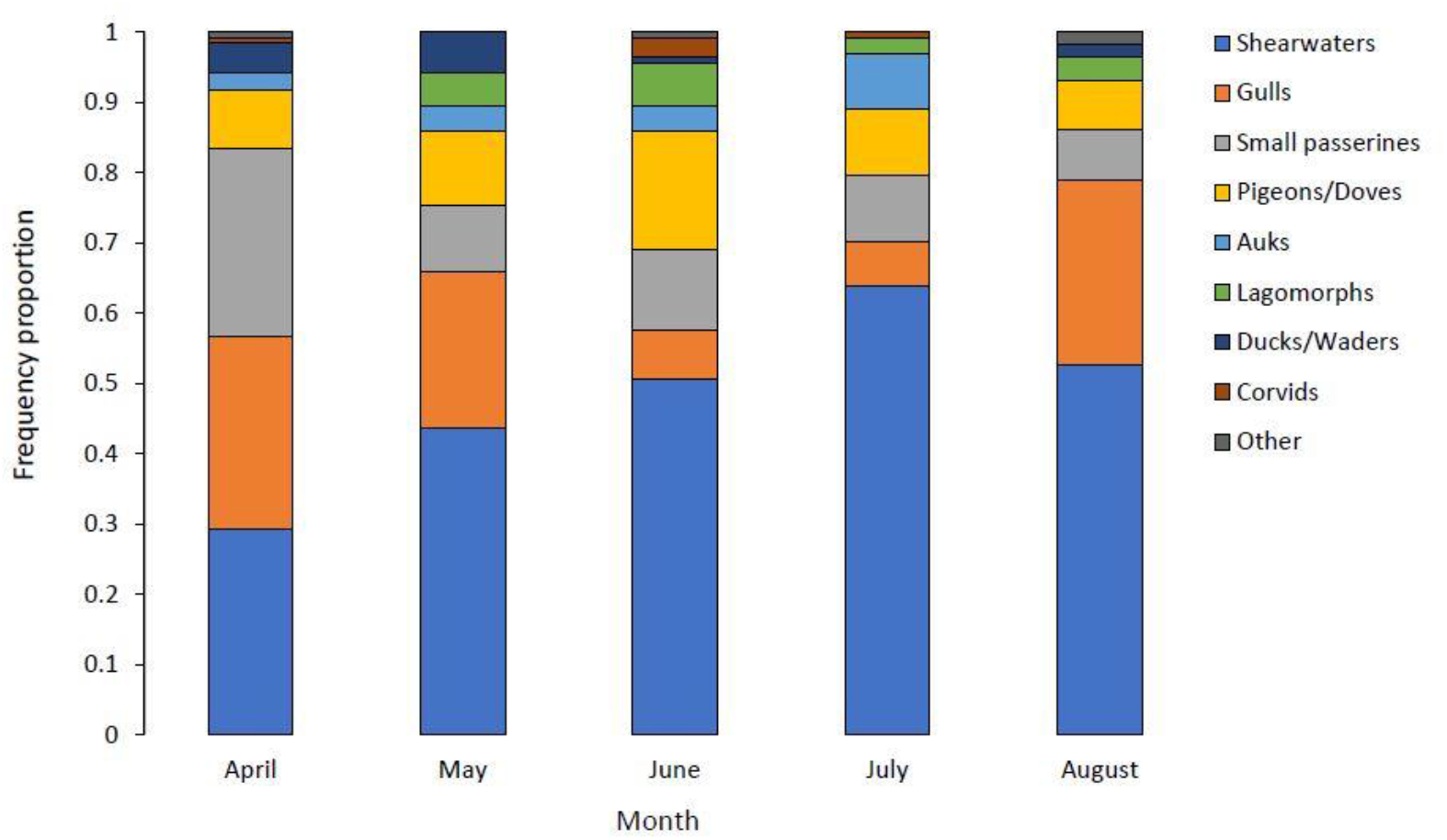
Temporal variation by month in frequency of prey groups predated by Peregrine Falcons breeding on Lundy from April-August 2014-2017.

The pair in territory L4/SW had the narrowest diet breadth and were the most specialized in terms of diet (*B*_*A*_ = 0.073), followed by the pairs in territory L1/NE (*B*_*A*_ = 0.116) and L5/W (*B*_*A*_ = 0.119) both with similar narrow diet breadth. Pairs at the remaining territories had relatively broader niche breadth and a more generalist diet (L2/E: *B*_*A*_ = 0.222; L3/S: *B*_*A*_ = 0.346). There was a strong inverse relationship between narrow dietary breadth and high percentage of Manx Shearwater in the diet of the territorial pairs (Fig. 5), with the percentage of Manx Shearwater in the diet explaining 87 % of the variation in dietary breadth (*F*_*1,3*_ = 20.2, *R*^2^ = 0.87, *p =* 0.02).

**Figure 5.**
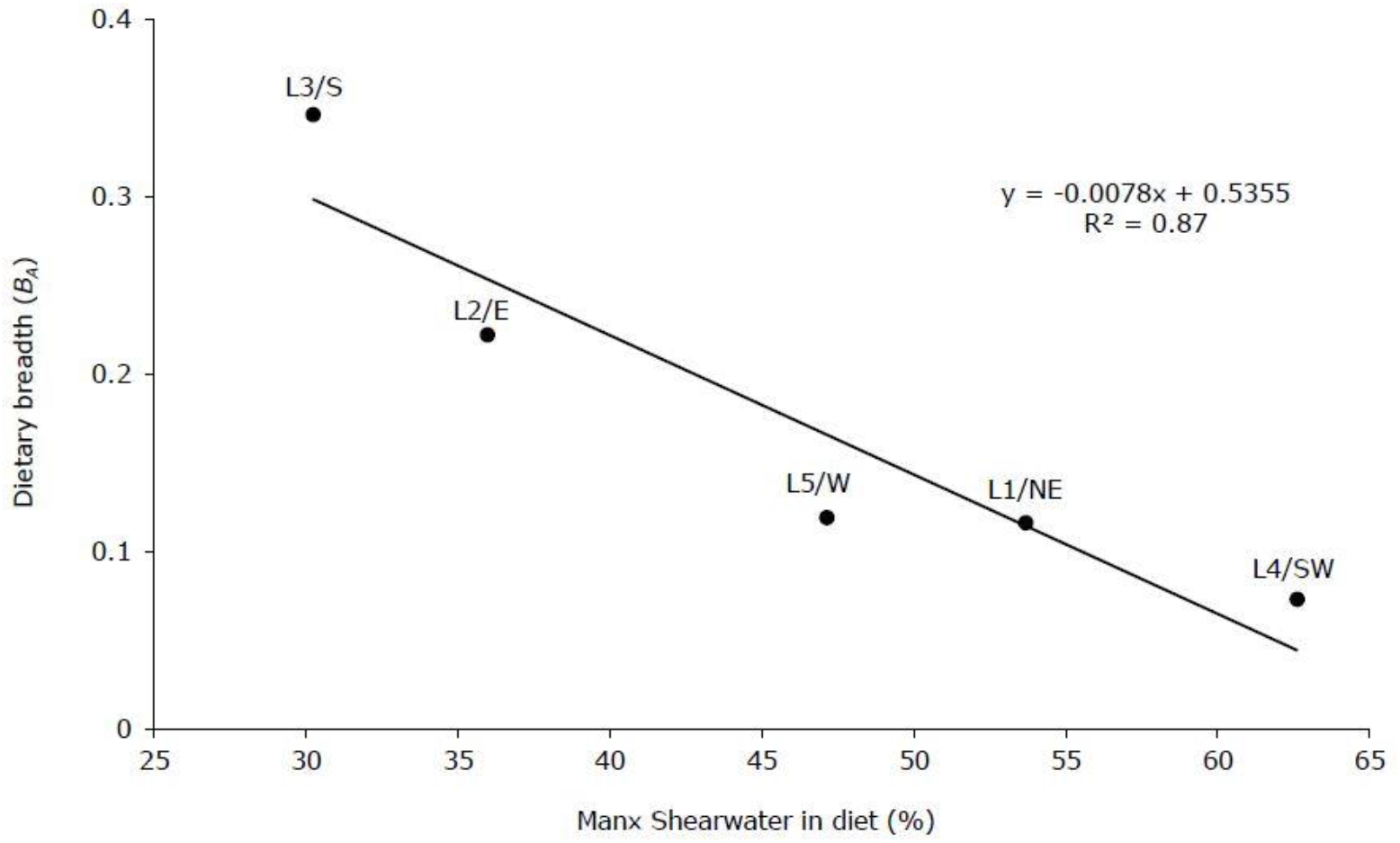
The relationship between Manx Shearwaters in total diet (%) and dietary breadth (*B*_*A*_). Each black point represents a breeding pair of Peregrine Falcons.

## Discussion

Our results demonstrate that specific breeding pairs of Peregrines on Lundy have a specialized diet, with Manx Shearwater the principal prey. This followed our general expectations given the marine location, though the finding that large prey such as Herring Gull and Rabbit were important food sources was unexpected. Prey mass was able to explain 16 % of the variation in predation frequency, with prey between 251-500 g the most common prey group. There was noticeable variation in spatial and temporal predation frequency, with the number of Manx Shearwater predated increasing substantially during June and July at three specific nest territories. Even so, all breeding pairs had narrow dietary breadth, with just three prey species accounting for ∼70 % of all prey taken. The trend of regular predation by Peregrines on specific prey types may have important implications for understanding IDS and its potential influence on insular raptor population structure.

Our results show the importance of Manx Shearwater in the diet of Peregrines breeding on Lundy. They were the most common prey species and group by both frequency and biomass (Fig. 1; Table S1). This finding is similar to other studies on insular Peregrine populations on offshore islands along the Pacific north west coast of North America. Here, the resident Peregrine sub-species *F*.*p. pealei*, has adapted to local prey availability, and specialises on feeding almost exclusively on Ancient Murrelets *Synthliboramphus antiquus* (Nelson 1990). Peregrine productivity and the number of breeding pairs fluctuates in accordance with seabird breeding density (Nelson & Myres, 1976; Nelson 2004). Understanding whether the number and distribution of the Manx Shearwater population on Lundy directly affects Peregrine breeding success is a key question to be determined from future research. Our results show that Shearwater predation frequency increases during June and July when food demand is highest from nestlings. Though there is no evidence that Peregrine breeding density is linked to the increase in the Manx Shearwater population, this abundant food supply could facilitate an increase in productivity for certain pairs in this Peregrine sub-population.

The mechanisms driving IDS in this insular Peregrine population are still unclear. Peregrines are highly territorial thus individuals could be restricted by territoriality between nesting pairs, limiting access to certain food resources (Ratcliffe 1993). Patchy habitats could be a key driver by providing different prey types to specific individuals, which then increase their foraging success and handling time by specialization (Sargeant 2007; Terraube *et al*. 2014). However, with such a dense population of Peregrines in a small area, intense intra-specific competition between breeding pairs for territory space, and therefore food, may be the strongest factor. Further research on home range size and foraging distances would be required to assess how Peregrines on Lundy defend territories and utilise habitats for these resources. Establishing if any broad-scale patterns of diet specialization exist for insular Peregrine populations across the British Isles may help inform conservation management for offshore islands in the region. Many of these islands hold globally important populations of seabirds which are often the main prey of Peregrines (Ratcliffe 1993). Therefore, considered management of seabird populations could indirectly benefit the predators that utilise them as a food resource.

Even though determining dietary habits from feeding remains is an established indirect method of assessing raptor diet (Marti *et al*. 2007), we recognise that indirect methods may over-represent large or brightly coloured prey species compared to small or darker prey remains that may go undetected (Ratcliffe 1993). Despite our thorough systematic sampling, we assume that some feeding areas were missed or inaccessible. Prey was observed being brought into nest ledges or feeding areas at territories L1/NE and L3/E that were inaccessible and out of sight. Thus, future dietary assessments of breeding season diet would benefit from direct methods using stable isotope analysis from tissue sampling of nestlings (Denhard *et al*. 2016; Horswill *et al*. 2016). It follows that the results presented here are only representative of what prey remains were observed and accessible.

Density compensation theory postulates that island species tend towards a generalist foraging strategy once released from competition with other species (Wright 1980). However, in some species, such as the Peregrine, this may be reversed and individuals within a given population may become diet specialists. Determining trophic relationships and relating those results to population dynamics in both predator and prey species is essential (Paine *et al*. 1990; Hipfner *et al*. 2011). This may establish how populations of avian apex predators are regulated on islands such as Lundy and how they interact in ecological community structures.

## Acknowledgements

LJS thanks the Birds of Lundy Fund, the Hawk and Owl Trust and the Lundy Field Society for providing financial assistance to conduct this research. We thank Dean Jones (Lundy Warden) who assisted with fieldwork and logistics.

## Appendix 1

**Table S1.** Frequency and biomass of all prey species taken by five breeding pairs of Peregrine Falcons on Lundy from April-August 2014-2017. Unit mass (g) taken from BTO (2016).

| Prey species | Frequency | Unit mass | Total biomass |
| --- | --- | --- | --- |
| Mallard *Anas platyrhynchos* | 4 (0.8) | 1210 | 4840 (2.1) |
| Eurasian Teal *Anas crecca* | 1 (0.2) | 323 | 323 (0.1) |
| Manx Shearwater *Puffinus puffinus* | 246 (47.3) | 383 | 94218 (40.8) |
| Common Kestrel *Falco tinnunculus* | 1 (0.2) | 220 | 220 (0.1) |
| Merlin *Falco columbarius* | 1 (0.2) | 202 | 202 (0.09) |
| Eurasian Woodcock *Scolopax rusticola* | 6 (1.2) | 291 | 1746 (0.8) |
| Common Snipe *Gallinago gallinago* | 1 (0.2) | 108 | 108 (0.05) |
| Eurasian Whimbrel *Numenius phaeopus* | 1 (0.2) | 451 | 451 (0.2) |
| Razorbill *Alca torda* | 7 (1.4) | 613 | 4291 (1.9) |
| Atlantic Puffin *Fratercula arctica* | 12 (2.3) | 387 | 4644 (2.0) |
| Black-headed Gull *Chroicocephalus ridibundus* | 2 (0.4) | 289 | 578 (0.3) |
| Lesser Black-backed Gull *Larus fuscus* | 3 (0.6) | 785 | 2355 (1.0) |
| European Herring Gull *Larus argentatus* | 72 (13.8) | 957 | 68904 (29.8) |
| Black-legged Kittiwake *Rissa tridactyla* | 7 (1.4) | 368 | 2576 (1.1) |
| Domestic Pigeon *Columba livia* | 45 (8.7) | 360 | 16200 (7.0) |
| Common Wood Pigeon *Columba palumbus* | 5 (1.0) | 507 | 2535 (1.1) |
| Eurasian Collared Dove *Streptopelia decaocto* | 3 (0.6) | 205 | 615 (0.3) |
| Common Cuckoo *Cuculus canorus* | 1 (0.2) | 114 | 114 (0.05) |
| Carrion Crow *Corvus corone* | 3 (0.6) | 509 | 1527 (0.7) |
| Eurasian Skylark *Alauda arvensis* | 11 (2.1) | 36 | 396 (0.2) |
| Barn Swallow *Hirundo rustica* | 5 (1.0) | 20 | 100 (0.04) |
| Long-tailed Tit *Aegithalos caudatus* | 1 (0.2) | 8 | 8 (0.01) |
| Common Starling *Sturnus vulgaris* | 2 (0.4) | 85 | 170 (0.07) |
| Common Blackbird *Turdus merula* | 1 (0.2) | 102 | 102 (0.04) |
| Song Thrush *Turdus philomelos* | 4 (0.8) | 75 | 300 (0.1) |
| Redwing *Turdus iliacus* | 2 (0.4) | 66 | 132 (0.06) |
| Mistle Thrush *Turdus viscivorus* | 2 (0.4) | 126 | 252 (0.1) |
| Common Chiffchaff *Phylloscopus collybita* | 2 (0.4) | 8 | 16 (0.01) |
| Willow Warbler *Phylloscopus trochilus* | 1 (0.2) | 9 | 9 (0.01) |
| European Robin *Erithacus rubecula* | 1 (0.2) | 19 | 19 (0.01) |
| Northern Wheatear *Oenanthe oenanthe* | 4 (0.8) | 28 | 112 (0.05) |
| House Sparrow *Passer domesticus* | 1 (0.2) | 27 | 27 (0.01) |
| Meadow Pipit *Anthus pratensis* | 15 (2.9) | 19 | 285 (0.1) |
| European Goldfinch *Carduelis carduelis* | 1 (0.2) | 16 | 16 (0.01) |
| Common Linnet *Carduelis cannabina* | 4 (0.8) | 19 | 76 (0.03) |
| *Alcidae* spp. | 1 (0.2) | 500 | 500 (0.2) |
| *Columbidae* spp. | 1 (0.2) | 357 | 357 (0.2) |
| *Corvidae* spp. | 2 (0.4) | 376 | 752 (0.3) |
| *Laridae* spp. | 7 (1.4) | 600 | 600 (0.3) |
| Unidentified Small Passerine | 15 (2.9) | 45 | 675 (0.3) |
| European Rabbit *Oryctolagus cuniculus* | 16 (3.1) | 1000 | 16000 (6.9) |
| Total | 520 (100) |  | 230951 (100) |

